# Non-Destructive Larval Genotyping of *Danio rerio* for Mitochondrial and Nuclear DNA Genetics

**DOI:** 10.1101/2025.04.30.651585

**Authors:** Kyler S. Mitra, Shannon R. Holmberg, Mireya Mota, Ankit Sabharwal, Stephen C. Ekker

## Abstract

The rapid advancement of nuclear and mitochondrial genomic editing tools has created an urgent need for efficient, non-lethal larval genotyping methods in zebrafish (*Danio rerio*) research. This study optimizes and validates a non-destructive proteinase K digestion method for mitochondrial and nuclear DNA genotyping while characterizing its impact on larval survival and gene expression. Using optimized protocol parameters, we demonstrate successful amplification of different mitochondrial and nuclear genetic loci with consistently high sensitivity. Molecular validation through PCR, restriction fragment length polymorphism analysis, and Sanger sequencing confirmed the specificity and reliability of the extracted DNA. The method successfully detected C-to-T base edits in the *mt-tl1* gene introduced using the FusX TALE Base editor system, demonstrating its applicability to gene editing studies. Both 48-well and optimized 96-well formats were used, enabling this approach to be deployed at scale. This optimized method enables researchers to correlate genotypes with phenotypes in longitudinal studies while maintaining specimen viability particularly valuable for investigating early-onset mitochondrial diseases and utilizes standard laboratory equipment and reagents, facilitating widespread adoption in zebrafish research while adhering to ethical principles in reducing animal mortality.

## Introduction

Zebrafish (*Danio rerio*) are the most widely used aquatic model system in scientific research, transcending their initial role in developmental biology to become instrumental in unraveling the genetic and biochemical intricacies of various cellular processes. This small vertebrate boasts an impressive array of advantages as a model organism, including rapid reproductive cycles, high offspring yield, external embryonic development, transparent embryos, swift organ formation, and straightforward husbandry requirements.^1–5^ The versatility of zebrafish has led to groundbreaking insights across a spectrum of biological processes and human diseases. From hematological and cardiovascular disorders to neurological conditions and cancer, zebrafish models have significantly contributed to our understanding of these complex systems.^6–8^ Zebrafish shine particularly bright in the realm of mitochondrial research. Their mitochondrial genome shares remarkable similarities with humans, with 100% gene synteny and approximately 65% sequence identity at the nucleotide level. This genetic kinship extends to codon usage, strand-specific nucleotide bias, and gene order, making zebrafish an ideal vertebrate model for studying human mitochondrial disorders.^9–11^ Since mitochondrial DNA (mtDNA) is extra-nuclearly segregated, it has different requirements for genotype to phenotype correlation. The complex genetics underlying mtDNA disorders, particularly their variable penetrance and diverse clinical presentations, can be attributed to the presence of either homoplasmy or heteroplasmy of mtDNA type within cells. The severity and progression of mitochondrial diseases are directly influenced by the ratio of mutated to normal (wild type) mtDNA within a cell.

Recent years have witnessed significant advancement in the mitochondrial DNA (mtDNA) editing toolbox such as DddA-derived cytosine base editors (DdCBEs) and TALE-linked deaminase (TALEDs) to engineer precise edits across different species enabling to model pathogenic mutations.^12–17^ Notably, DddA-derived cytosine base editors (DdCBEs) have achieved editing efficiencies at targeted mtDNA loci, enabling accurate modeling of pathogenic mutations. Additionally, engineered variants such as DddA6 and DddA11 have expanded the editing scope by targeting previously inaccessible sequence contexts, while monomeric DdCBE (mDdCBE) constructs facilitate broader applicability and simplified delivery.^13,18^ These developments collectively enhance the utility of zebrafish as a robust vertebrate model for mitochondrial genetic research. The advent of these mitochondrial genomic editing tools underscores the critical need for rapid and efficient larval genotyping in zebrafish research. Traditional approaches to live larval genotyping have relied on microsurgical fin biopsy performed in water droplets,^19^ while alternative methods involve the complete lysis of embryos for cohort genotype determination^20^ or an automated genotyping device.^21^ However, these conventional techniques present significant limitations: fin microsurgery is labor-intensive and frequently lethal, while embryonic lysis precludes the survival of genotyped larvae for subsequent analysis or breeding. Non-destructive enzyme genotyping presents a scalable solution that overcomes these constraints by employing standard laboratory reagents and materials. This technique facilitates DNA extraction through controlled proteinase K (ProK) digestion of epithelial cells, yielding genetic material that can be PCR amplified while preserving larval viability.^22^ This protocol enables greater efficiency in husbandry and high-throughput assays, enabling genotype determination within days of fertilization. Notably, this non-lethal approach proves particularly valuable for investigating early-onset lethal phenotypes, where rapid genotyping of living specimens is essential for understanding disease progression.

In this study, we employed and refined a non-destructive genotyping protocol for the mitochondrial and nuclear genome and characterized the impact of these assays on larvae through survival and transcriptomics data. During testing and troubleshooting of this non-destructive assay, we identified both advantages and disadvantages of the proteinase K method. Additionally, we demonstrated successful amplification of mitochondrially encoded genes alongside nuclear genes. Finally, modifications to the previously published enzyme protocol have enabled stronger survival.

## Materials and Methods

### Zebrafish handling and husbandry

All adult zebrafish and embryos were maintained according to the guidelines established by Mayo Clinic Institutional Animal Care and Use Committee (IACUC number: A34513-13-R16) and The University of Texas at Austin Institutional Animal Care and Use Committee (IACUC number: AUP-2023-00200).

### Proteinase K-based larval genotyping

Wild-type (WT) Mayo Recessive Free (MRF) zebrafish and mutant transgenic lines, *mt-tl1*^14^ were used to generate larvae for non-destructive genotyping.^22^ Three days post-fertilization (dpf) larvae were rinsed three times in embryo media followed by three washes in tricaine buffer (30 mM Tris-HCl in 0.4% tricaine solution). Using a p200 pipette tip trimmed to accommodate larval width, individual larvae were aspirated in 40 µL of ProK genotyping buffer (25 µg/mL Proteinase K (catalog no.: EO0491, Thermo Fisher Scientific, USA) in tricaine buffer) and placed in 96-well (catalog no.: 3455, Thermo Fisher Scientific, USA) and 48-well plates (catalog no.: CC7672-7548, CytoOne). For 48-well plates, droplets were dispensed centrally within each well to minimize contact between larvae and well walls during agitation. Proteinase K concentration and incubation time can be adjusted based on larval age and condition to optimize survival rates. The plate was incubated in a heated shaker (catalog no.: VORTEMP 56, Labnet International, Inc., USA) at 37°C with 500 rpm shaking for 20 minutes. After incubation, 30 µL of the liquid around the fish was carefully pipetted (avoiding larval aspiration) into 15 µL of lysis buffer (10 mM Tris-HCl, pH 8.0, 50 mM KCl, 0.3% Tween-20, 0.3% NP-40, 1 mM EDTA) in 0.2 mL strip tubes and pipetted up and down ∼10 times. The solution was then incubated at 98°C for 5 minutes to extract DNA. Following media collection, wells were quickly replenished with embryo media using a Pasteur pipette, and larvae were transferred to 24-well plates containing fresh embryo media.

To prevent tail damage during the shaking procedure, plates can be pre-treated with a protein-rich blocking solution. If extensive tail damage is observed within shaken larvae, plates can be incubated with 5% non-fat dry milk (NFDM) solution prepared in embryo media for 5 minutes, followed by three washes with embryo media prior to genotyping (Fig. 1A).

**Figure 1:**
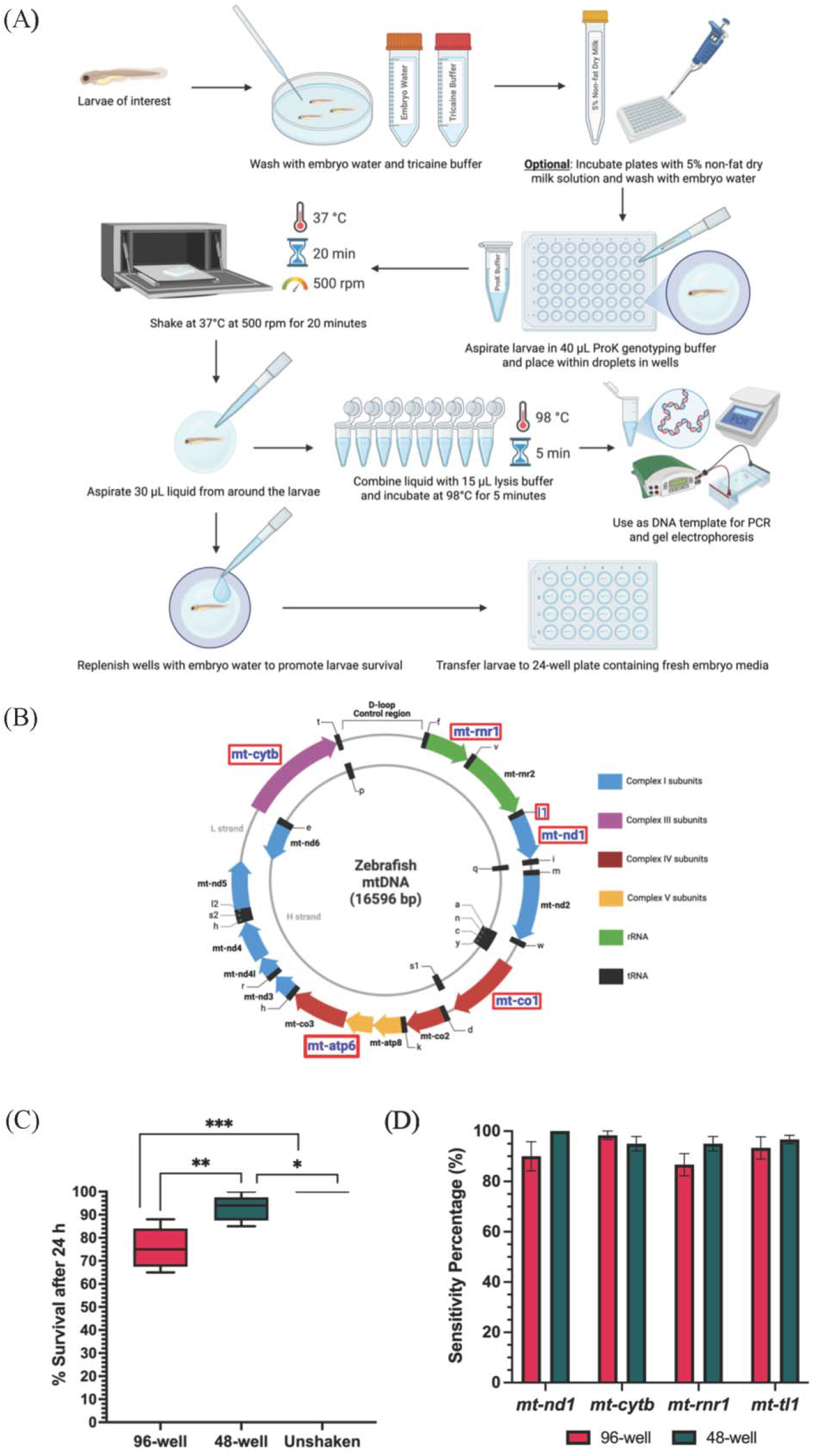
Non-Destructive Enzyme Genotyping Workflow and Validation of Survival and Sensitivity. **(A)** Schematic of non-destructive enzyme genotyping protocol. **(B)** Schematic of the zebrafish mitochondrial genome showing genes targeted for non-destructive genotyping (highlighted in red boxes) **(C)** Box-and-whisker plot comparing survival percentages of larvae 24-hour post genotyping in 96-well, 48-well, and unshaken control groups. Whiskers represent maximum and minimum values. p-Values were determined by Student’s t-test (*p□<□0.05; **p□<□0.01; ***p<0.001). (N=100 technical replicates across 5 experiments). **(D)** Sensitivity of the protocol is represented by the percentage of successfully amplified samples across four mitochondrial genes (N=60 technical replicates across 3 experiments). No statistically significant difference was observed between the well formats with respect to the amplicon sensitivity. Error bars are represented as standard error of mean. Figure 1 A and B was created using Biorender.com.

### Survival assay

Larvae collected after enzymatic genotyping were transferred into individual wells of a 24-well plate. Unshaken age-matched zebrafish larvae obtained from the same clutch were used as controls. Live counts for larvae from each group were recorded at 24 hours post-genotyping, and morphological defects were examined pre- and post-procedure.

### PCR amplification and Restriction Fragment Length Polymorphism for mitochondrial genetic loci

To investigate the sensitivity of the protocol for mitochondrial DNA genotyping, we prioritized protein-coding genes from each of the respiratory chain complex, transfer RNA and ribosomal RNA genes. We targeted six mitochondrial genetic loci with distinct functional roles. Four of them were protein coding genes: mitochondrially encoded NADH:ubiquinone oxidoreductase core subunit 1 (*mt-nd1*), mitochondrially encoded cytochrome b (*mt-cytb*), mitochondrially encoded ATP synthase membrane subunit 6 (*mt-atp6*) and mitochondrially encoded cytochrome c oxidase subunit 1 (*mt-co1*). Remaining two were non-coding genes: mitochondrially encoded tRNA-Leu (*mt-tl1*) and mitochondrially encoded 12S rRNA (*mt-rnr1*) (Fig. 1B). PCR amplification was performed for each of the six genes using their respective primers mentioned in Supplementary Table 1. PCR consisted of 10□μL of 2X OneTaq master mix with standard buffer (NEB, USA), 1□μL of 10□μM forward primer, 1□μL of 10□μM reverse primer and 8□μL of template. Samples were run in a thermal cycler with the following PCR conditions: (1) 95°C, 4□min; (2) 95°C, 30 sec; (3) 60°C, 30□sec; (4) 72°C, 30□s; (5) repeat steps 2-4 for 30□cycles□; (6) 72°C, 4□min; and (7) 12°C, hold. PCR amplicons were run on 1% agarose gel and analyzed.

Five to eight microliters of the PCR amplicon (depending on intensity of the amplicon on the gel) were used as the substrate for restriction fragment length polymorphism (RFLP) in 20 µL reaction solution (2 µL of the 10x corresponding buffer and 5 U of restriction enzyme and remaining volume with nuclease-free water). PCR amplicons for genetic loci *mt-nd1, mt-cytb, mt-tl1*, and *mt-rnr1* were surveyed for restriction digestion by SpeI, AvaI, SacI and HindIII, respectively. A subset of the PCR amplicons were then submitted to Sanger sequencing at GENEWIZ (Azenta, USA) to confirm the sequence specificity. Chromatograms files (.ab1) were then uploaded to the EditR^23^ web server to estimate the heteroplasmy in the mtDNA mutants.

### PCR amplification of nuclear genetic loci

To investigate the sensitivity of the protocol for nuclear gene genotyping, we prioritized protein-coding nuclear genes with roles in mitochondrial function and signaling. The three nuclear loci were: mitochondrial antiviral signaling (*mavs*), cytochrome c oxidase subunit 7B (*cox7b*), and leucine-rich pentatricopeptide repeat-containing (*lrpprc*). PCR amplification was performed for each of these genes using their respective primers listed in Supplementary Table 1. PCR reactions consisted of 10□μL of 2X OneTaq master mix with standard buffer (NEB, USA), 1□μL of 10□μM forward primer, 1□μL of 10□μM reverse primer, and 8□μL of template DNA. PCR amplification was carried out under the following conditions: (1) 95°C, 4□min; (2) 95°C, 30□s; (3) annealing at 58°C for *cox7b* and *lrpprc*, or 54°C for *mavs*, 30□s; (4) 72°C, 30□s for *mavs* and *lrpprc*; and 2 min for *cox7b* (5) repeat steps 2–4 for 34 cycles; (6) 72°C, 4□min; (7) hold at 12°C. PCR amplicons were run on 1% agarose gel and analyzed. A subset of the PCR products was submitted to Sanger sequencing at GENEWIZ (Azenta, USA) to confirm sequence specificity.

### RNA extraction and sample preparation

To assess for any differential transcriptional signatures in the proteinase K-shaken larvae to those unshaken, we performed whole body RNA sequencing for the two groups. Adult zebrafish were incrossed and non-destructive genotyping was performed on larvae aged 3 dpf. Larvae were subjected to the protocol for the non-destructive enzymatic proteinase K conditions and were then allowed to recover in embryo water until 6 dpf. Six larvae were pooled, and RNA isolation was performed using the Qiagen RNeasy Mini kit as per manufacturer’s instructions. An additional DNAse treatment step was performed before the final wash and elution. RNA concentration was measured using a spectrophotometer.

### RNA sequencing and analyses

RNA library preparations and sequencing reactions were performed at GENEWIZ (Azenta USA). For transcriptome sequencing, RNA was isolated by pooling six individual larvae from each condition. RNA quantification was performed using the Qubit 2.0 Fluorometer (Thermo Fisher Scientific, USA), and RNA integrity was assessed using the Agilent TapeStation (Agilent Technologies, USA). RNA sequencing libraries were constructed following the NEBNext Ultra II RNA Library Prep Kit for Illumina protocol (NEB, USA). Briefly, mRNA was enriched using oligo(dT) beads, fragmented for 15 minutes at 94°C, and then reverse-transcribed into cDNA. The cDNA was end-repaired, adenylated at the 3’ ends, and ligated with universal adapters, followed by index addition and PCR-based library amplification. The sequencing library was validated on the Agilent TapeStation (Agilent Technologies, USA) and quantified using both the

Qubit 2.0 Fluorometer and quantitative PCR (KAPA Biosystems, USA). The sequencing libraries were then multiplexed, clustered onto a flow cell, and loaded onto the Illumina HiSeqX instrument according to the manufacturer’s instructions. Sequencing was performed using a 2×150 bp Paired-End (PE) configuration. Image analysis and base calling were carried out with HiSeq Control Software (HCS). The raw sequence data (.bcl files) generated from the Illumina HiSeqX platform were converted into fastq files and demultiplexed using Illumina’s bcl2fastq 2.20 software with one mismatch allowed for index sequence identification. The raw fastq data was uploaded to the Basepair cloud bioinformatics platform (https://www.basepairtech.com/). Filtered reads were aligned to the zebrafish reference genome (Zv11 version) using STAR. Raw and normalized gene and transcript counts were calculated. Differential expression analysis was performed using DESeq2^24^, with a threshold of p-adjusted <0.05 for identifying differentially expressed genes between the genotyped and unshaken control larvae.

## Results

### Survival rates of enzyme-genotyped zebrafish larvae support high-throughput screening

Non-destructive enzyme genotyping of zebrafish larvae demonstrates high survival rates and compatibility with high-throughput screening. Larvae shaken in 96-well plates without milk pretreatment had a median survival rate of 75%, while 48-well larvae had a median of 94%, compared to 100% in unshaken controls (Fig. 1C). Most larval deaths occurred within 24 hours post-shaking, primarily due to tail fin damage, with no observed differences in survival between undamaged shaken and unshaken fish after this period. Key survival-enhancing steps included rapidly replacing embryo water after DNA collection and promptly removing larvae from the heated shaker. These parameters can be further adapted to accommodate different larval ages and sensitive phenotypes. Importantly, no morphological changes were observed in surviving genotyped larvae compared to unshaken controls, suggesting the method’s minimal physiological impact.

### Mitigating characteristic tail fin damage among ProK-genotyped larvae via 48-well plates and milk solution pretreatment

Our initial attempts using 96-well plates for ProK genotyping resulted in a subset of larvae exhibiting fatal damage, including partial to complete tail fin tearing. We attributed this damage pattern to tail fin adhesion to the well walls during heated shaking, exacerbated by ProK-induced weakening of fin structural integrity. We determined this damage pattern likely resulted from tail fin adhesion to well walls during heated shaking, exacerbated by ProK-induced weakening of fin structural integrity. Two effective mitigation strategies were thus developed. First, utilization of 48-well plates enabled placement of larvae in centered droplets without wall contact, significantly increasing survival and reducing tail fin damage. However, the efficacy of this approach may vary depending on the strength of the hydrophobic coating applied by different manufacturers to maintain protective droplet formation. Alternatively, pretreating 96-well plates with 5% non-fat dry milk (NFDM) solution created a protective proteinaceous film that reduced larval tail fin adhesion during shaking (Fig. 1A). We observed decreased tail fin damage and improved survival rates among genotyped larvae in the 96-well plate treated with 5% NFDM in embryo media (Supplementary Fig. 1).

### High sensitivity of amplification across diverse mitochondrial genetic loci

Non-destructive enzyme genotyping demonstrates high amplification sensitivity across diverse mitochondrial genes encoding critical cellular machinery. Amplification sensitivity was assessed for *mt-nd1, mt-cytb, mt-rnr1*, and *mt-tl1* by calculating the percentage of successfully amplified samples across these genes. For each gene-well combination, we conducted three experimental trials, with each trial containing 20 larvae. Thus, we analyzed a total of 60 larvae for each gene-well combination, resulting in 120 total samples per gene across both formats. Consistently high amplification rates were observed across all four genes in both formats. All experimental sets exhibited over 80% sensitivity (Fig. 1D), demonstrating robust performance of this non-destructive genotyping method across diverse mitochondrial genes.

### Specificity of mitochondrial DNA amplification verified by PCR and RFLP analysis

We validated the specificity of our non-destructive enzyme genotyping method through comprehensive molecular analysis of four mitochondrial genes. PCR amplification and restriction fragment length polymorphism (RFLP) analysis were performed on wild-type zebrafish larval DNA samples from 96-well and 48-well plates, with all samples referenced against Quick-Load Purple 50 bp DNA Ladder (NEB, USA). For *mt-nd1*, we observed a 350 bp PCR amplicon with SpeI restriction digest yielding 236 bp and 114 bp fragments (Fig. 2A, B). The *mt-cytb* gene generated a 460 bp PCR amplicon with AvaI restriction digest yielding 317 bp and 143 bp fragments (Fig. 2D, E). Amplification of *mt-rnr1* showed a 395 bp PCR amplicon with HindIII restriction digest producing 253 bp and 142 bp fragments (Fig. 2G, H). The *mt-tl1* gene produced a 363 bp PCR product (Fig. 3A, B) with SacI restriction digest creating 186 bp and 177 bp fragments. PCR amplification of *mt-atp6* produced a 369 bp amplicon (Supplementary Fig. 2A), while *mt-co1* generated a 494 bp PCR product (Supplementary Fig. 2B). Sanger sequencing confirmed the presence of restriction enzyme sites for the first four genes (Fig. 2C, F, I, 3E), and all samples selected for gel visualization represented the median amplification range within their respective cohorts.

**Figure 2:**
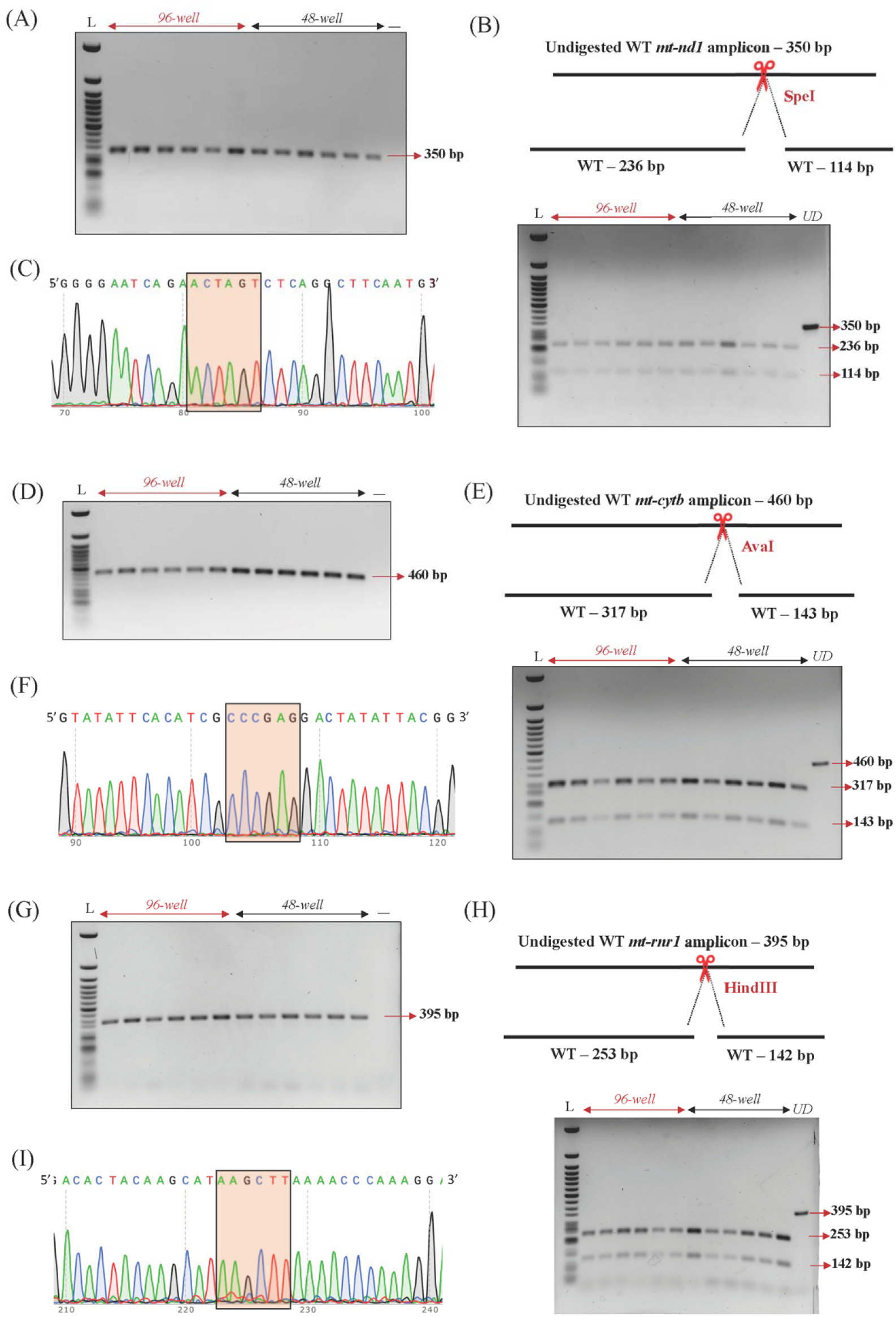
Specificity of Mitochondrial DNA Amplification. **(A)** Gel electrophoresis of the mitochondrial gene *mt-nd1*, showing amplification of the 350 bp band. **(B)** Screening of *mt-nd1* amplicon by RFLP. PCR amplicons from wild type larvae show the presence of uncut product (350□bp) and expected 236 and 114□bp digested bands. UD = undigested PCR product. **(C)** Representative chromatogram of the *mt-nd1* amplicon depicting the restriction enzyme recognition site. **(D)** Agarose gel depicting amplification of the mitochondrial gene *mt-cytb*, with the expected 460□bp fragment. **(E)** RFLP analysis of the *mt-cytb* PCR product, showing the intact 460 bp band and the predicted 317 and 143□bp digestion fragments. UD = undigested PCR product. **(F)** Representative electropherogram of the *mt-cytb* amplicon, with the restriction enzyme recognition site indicated. **(G)** Visualization of *mt-rnr1* amplification, showing a discrete 395□bp band. **(H)** Restriction digest of the *mt-rnr1* PCR product, displaying both the undigested 395□bp fragment and the expected 253 and 142□bp cleavage products. UD = undigested PCR product. **(I)** Sanger sequencing trace of the *mt*□*rnr1* amplicon, highlighting the enzyme cut site.

**Figure 3:**
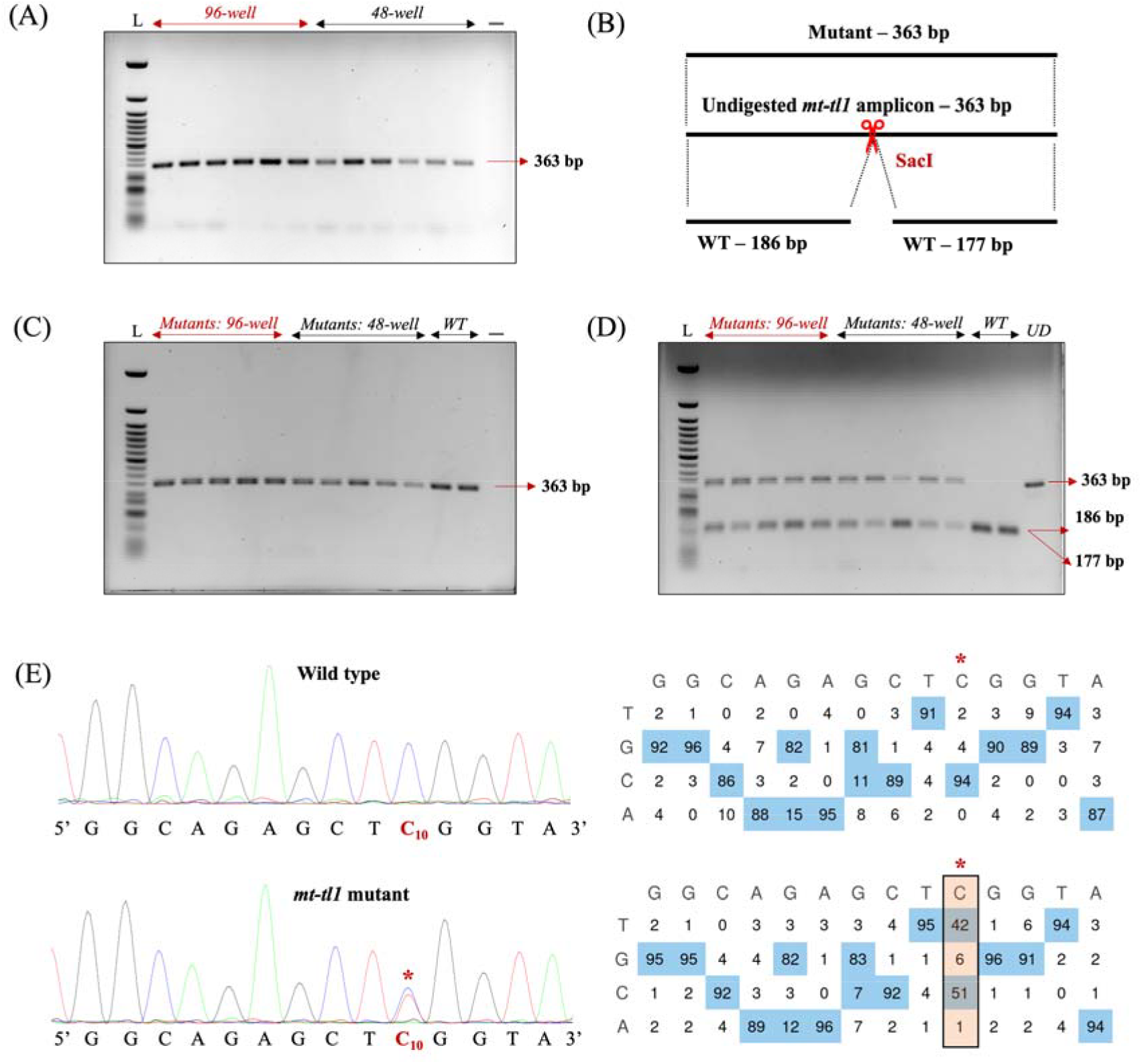
Screening for mtDNA mutants using non-destructive genotyping in *mt-tl1* locus. **(A)** PCR amplification of the *mt*□*tl1* loci from wild□type larvae resolved on a 1% agarose gel to confirm product integrity. **(B)** Schematic of the SacI recognition site within the amplicon (363 bp) and the expected fragment sizes (186 and 177 bp) for both unedited and edited alleles. **(C)** PCR products from *mt-tl1* mutants are shown on an agarose gel, demonstrating successful amplification. **(D)** RFLP analysis reveals that C□to□T conversion at the target cytosine leading to loss of SacI recognition site: control larvae yield the expected 186□bp and 177□bp fragments, whereas edited samples display a single undigested band (each lane represents individual sample DNA; UD = undigested PCR product). **(E)** Sanger sequencing chromatograms from control and mutant larvae. Asterisk (*) denotes the site of edit with corresponding editing percentage (C-to-T). Chromatograms and editing table plot were obtained using EditR.

### Genotyping of mtDNA mutant

We demonstrated the utility of our non-destructive genotyping approach for gene editing by introducing a C-to-T base substitution in zebrafish *mt*□*tl1* mutants.^14^ Three dpf *mt-tl1* zebrafish mutants were genotyped non-destructively, enabling subsequent longitudinal studies (Fig. 3C). The engineered C-to-T transition at position m.C3744 (C10 on the forward strand) resulted in the loss of the SacI restriction site, thereby facilitating the identification of larvae with mtDNA variations. RFLP analysis demonstrated the loss of the SacI restriction site in mutant larvae, with digested products of 186 bp and 177 bp, and an undigested product of 363 bp. Wild-type controls from both 96-well and 48-well cohorts showed only digested products of 186 bp and 177 bp (Fig. 3D). RFLP analysis was confirmed by Sanger sequencing where sequencing files (.ab1) were assessed using EditR^23^ software to quantify the level of editing in the mutant larvae. Sanger sequencing confirmed the C-to-T edit, revealing over 40% mtDNA mutant heteroplasmy levels at the target locus (Fig. 3E).

### Amplification across different nuclear genetic loci

To expand the utility of the optimized protocol, we adopted non-destructive genotyping to amplify different nuclear genetic loci. We observed successful amplification of the nuclear genes *mavs* (Fig. 4A, B), *cox7b* (Fig. 4C, D) and *lrpprc* (Fig. 4E, F) by agarose gel electrophoresis (Fig. 4). PCR amplicons for *mavs* and *lrpprc* were referenced against Quick-Load Purple 50 bp DNA Ladder (NEB, USA), and for *cox7b*, amplicons were referenced against HyperLadder 1kb (Meridian Bioscience, USA) Sanger sequencing confirmed that PCR products corresponded to the intended nuclear gene loci. This demonstrated that these optimizations can be employed for nuclear DNA genotyping as well.

**Figure 4:**
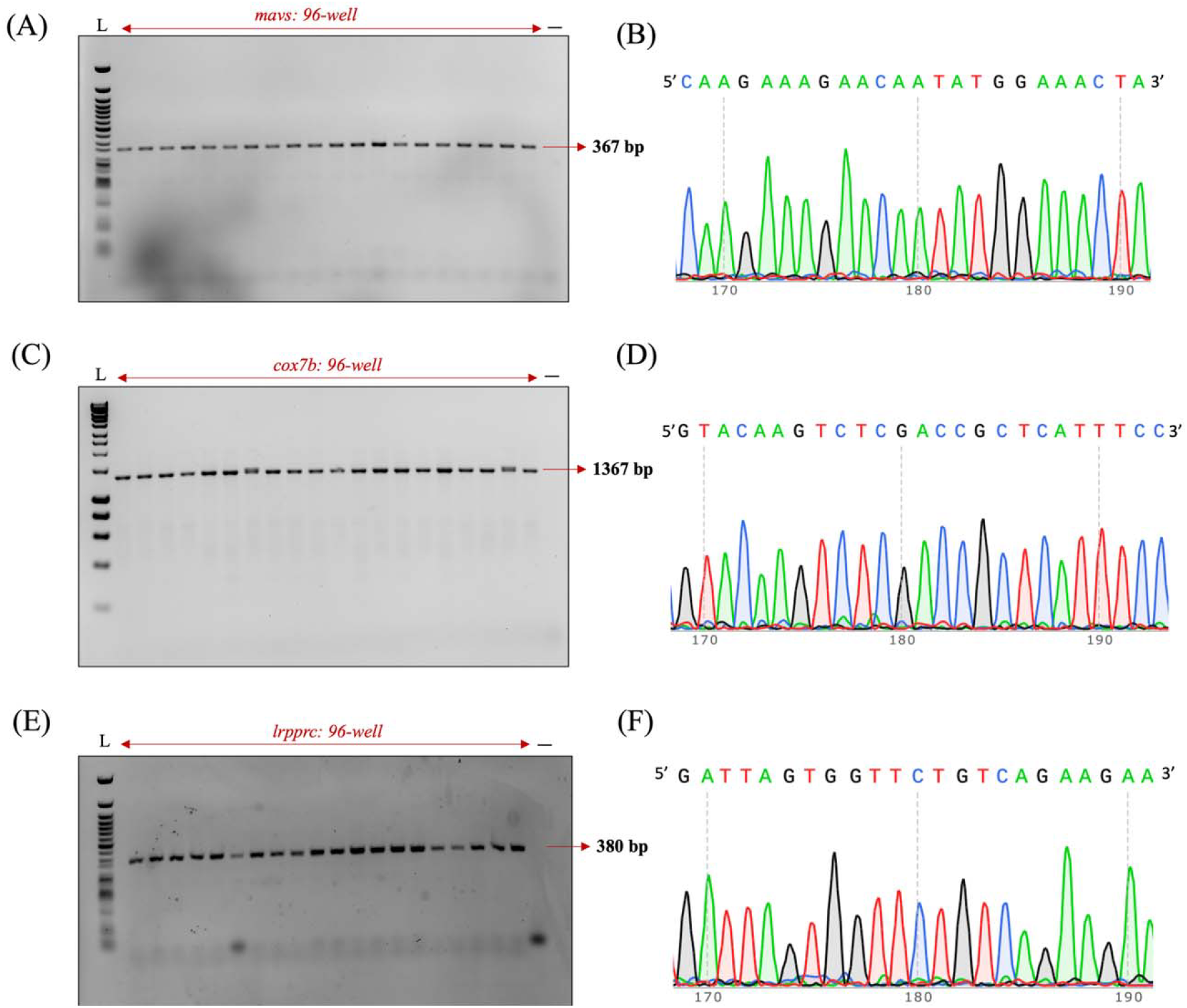
Specificity of Nuclear DNA Amplification. **(A–F)** Gel electrophoresis demonstrates successful PCR amplification for each nuclear locus: *mavs* **(A)** yields a distinct 367 bp product; *cox7b* **(C)** a 1367 bp product; and *lrpprc* **(E)** a 380 bp product, corresponding to expected amplicon size for each locus. **(B, D, F)** Representative Sanger sequencing chromatograms for the *mavs* **(B)**, *cox7b* **(D)**, and *lrpprc* **(F)** amplicons show high-quality, clearly resolved peaks, confirming the fidelity of PCR amplification and enabling downstream sequence analysis.

### Transcriptomic signature in non-destructively genotyped zebrafish larvae

To examine the genome-wide transcriptional changes in zebrafish larvae following non-destructive genotyping, we performed strand-specific paired-end RNA sequencing to identify a candidate differential expression profile. Sequence data were processed using the DESeq2^24^ pipeline on the Basepair cloud bioinformatics platform (https://www.basepairtech.com/). For both Proteinase K-shaken larvae and wild-type controls, we generated between 34 and 39 million reads. Reads below a Phred score of Q30 were discarded, and the remaining high-quality reads were aligned to the Zv11 zebrafish reference genome, with approximately 78% mapping uniquely. These uniquely mapped reads were then assembled to quantify transcript levels across the genome. Upon applying the criteria of a log□ fold change of ≥1 or ≤–1 with an adjusted p-value < 0.05 in the dataset (Supplementary Fig. 3), zero downregulated genes were observed. We identified seven annotated genes and six uncharacterized genes to be significantly upregulated in the 96-well genotyping treatment as compared to unshaken controls (Supplementary Fig. 3 and Supplementary File 1).

## Discussion

In this study, we optimized and validated a non-destructive larval genotyping method for efficient molecular analysis of zebrafish mitochondrial and nuclear DNA while maintaining high survival rates. Building upon the non-destructive genotyping protocol originally established by Zhang et al.,^22^ we demonstrated its utility specifically for mitochondrial DNA genetics and adapted it for high-throughput applications, allowing simultaneous processing of up to 96 larvae. Our approach addresses a significant challenge in zebrafish research—the ability to collect sufficient DNA for reliable genotyping from larvae without sacrificing the animal or compromising its development, which is inherently important for correlating genotype to phenotype in mitochondrial genetics. The optimized protocol achieves high amplification sensitivity across diverse mitochondrial and nuclear genes while preserving larval viability.

The protocol provides a balanced approach for mtDNA genotyping that minimizes stress to the larvae. Notably, the high survival rates observed, particularly in 48-well plates and 96-well with 5% NFDM, demonstrated the method’s suitability for high-throughput screening applications. Our technical improvements, particularly the implementation of 48-well plates and milk pretreatment steps, significantly mitigated tail fin damage—a principal cause of larval mortality in our earlier approaches. These modifications addressed the critical issue of tail fins adhering to well walls during heated shaking, which when combined with ProK-induced weakening of fin structural integrity, frequently resulted in fatal damage including torn tail fins. The 48-well plates allowed for centered droplet placement that prevented wall contact, while the milk pretreatment created a proteinaceous coating that reduced the adhesive properties of the well surfaces, preventing tail fins from sticking to the walls during agitation. Together, these adjustments substantially improved survival rates, enabling effective non-destructive genotyping without compromising experimental viability. The absence of morphological changes in surviving larvae, supported by our mRNA transcriptomics data showing no significant changes in the mitochondrial gene expression, further confirmed the minimal physiological impact of the protocol on mitochondrial homeostasis. These findings are particularly valuable for longitudinal studies and subsequent phenotypic analyses that require the same individual to be monitored over time. Several technical considerations emerged during out optimization process. We found that rapid replenishment of embryo water post-DNA collection and prompt removal from the heated shaker significantly improved larvae survival. These insights highlight the importance of minimizing exposure to Proteinase K solutions and elevated temperatures. Additionally, our observation that most larval mortality occurred within 24 hours of shaking due to tail fin damage suggests areas for further protocol refinement. The DNA collected through this non-destructive method was found to be stable for long periods comparable to conventionally collected fin clip DNA, with successful amplification and sequencing more than 6 months post-collection.

Our results demonstrated consistent amplification success (>80% sensitivity) across different mitochondrial DNA and nuclear DNA genetic loci. The robust performance across these diverse genetic loci suggested that our method could reliably detect a range of mitochondrial and nuclear sequences, which is particularly important for studies examining mitochondrial and nuclear genetics. The molecular validation through PCR, RFLP analysis, and Sanger sequencing confirmed the specificity and reliability of the genotyping method. The precise restriction enzyme digestion patterns obtained for each mitochondrial gene demonstrated the specificity of the extracted DNA. This methodological validation is crucial for ensuring the accuracy of downstream genetic analyses and interpretations. Additionally, we successfully amplified two other important mitochondrial genes—*mt-atp6* and *mt-co1*—conducting only PCR amplification for these loci with strong amplification throughout. Significantly, we successfully applied our non-destructive genotyping method to detect gene editing changes in the *mt-tl1* mutants established earlier using the FusX TALE Base Editor system.^14^ The detection of C-to-T base edits with over 40% mtDNA mutant heteroplasmy levels underscored the protocol’s sensitivity for identifying genetic modifications. This capacity is particularly pertinent for mitochondrial disorders, where early detection of heteroplasmic variations is essential for elucidating pathogenic mechanisms.

The RNA-seq analysis revealed a small number of significantly upregulated genes, several of which are known regulators of circadian rhythm and transcriptional control. Notably, period circadian clock 1a (*per1a*), nuclear receptor subfamily 1 group d member 1 (*nr1d1*), and aryl hydrocarbon receptor nuclear translocator-like protein 2 (*arntl2*) were among the most strongly induced. Their coordinated overexpression suggests that the experimental condition may directly influence circadian rhythm regulation.^25^ In addition to these well-characterized regulators, the dataset highlights several uncharacterized or poorly annotated transcripts, including LOC101886224, LOC100535095, and LOC101883914, which showed moderate to strong upregulation. Interestingly, granulin 2 (*grn2)* and granulin antisense (*grnas*) was also significantly upregulated. Grn2 protein belong to family of proteins known as granulins that play a role in cell proliferation, growth and inflammation.^26^ However, we didn’t find any mitochondrial resident nuclear encoded genes or mitochondrial DNA encoded genes to be differentially expressed in the genotyped larvae. These results highlight an important point to consider that while the method enables efficient genotyping of live zebrafish for longitudinal research, it can transiently alter gene expression profiles. This effect has the potential to confound downstream results, especially if the genes or pathways altered by genotyping overlap with those under investigation. Laboratories should consider whether genotyping could influence cellular homeostasis or other key processes central to their studies and adopt controls accordingly during experimental analysis.

Another important consideration for downstream experiments following genotyping is the incubation temperature. Incubation at 37□°C may transiently activate heat shock response pathways, particularly in transgenic lines carrying heat shock–inducible promoters. Although our RNA-seq analysis did not detect heat shock–related gene expression changes, we cannot exclude the possibility of transient stress-related transcriptional responses immediately after treatment.

By preserving larval integrity, this method enables longitudinal analyses that are crucial for determining the germline inheritance of mitochondrial variations, an aspect that remains underexplored in current zebrafish models. Additionally, the ability to correlate specific mtDNA mutations with metabolic dysfunction and disease progression opens avenues for more detailed phenotypic and metabolic assays. This continuous tracking across developmental stages not only enhances the robustness of genotype–phenotype correlations but also paves the way for translational applications in modeling mitochondrial diseases with early onset lethality. Moreover, the high-throughput nature of the protocol facilitates the screening of large sample sizes, enhancing statistical power when investigating rare mitochondrial variants and their associated phenotypes.

In conclusion, the optimization of the non-destructive enzyme genotyping method provides a valuable tool for zebrafish researchers, enabling efficient molecular characterization of both nuclear and mitochondrial genomes while maintaining animal viability. This approach not only enhances the ethical aspects of zebrafish research by reducing animal mortality but also expands the experimental possibilities for developmental, genetic, and disease modeling studies, particularly for conditions with early onset lethality.

## Acknowledgments

The authors would like to thank the staff of the Mayo Clinic Zebrafish Facility and the University of Texas at Austin Zebrafish Facility. We would also like to acknowledge Dr. Karl J. Clark, Texas A&M University, Texas, USA for the valuable suggestions and feedback.

## Authors’ Contributions

The idea was conceived by A.S and S.C.E. The article was written by K.S.M, S.R.H, A.S., and S.C.E. Experiments were executed by K.S.M., S.R.H., M.M., and A.S. with experimental guidance from S.C.E. Data analysis was completed by K.S.M, S.R.H., and A.S. The article was reviewed and edited by K.S.M, A.S, and S.C.E.

## Data Availability

Raw sequencing data has been uploaded on NCBI SRA (SRA ID: PRJNA1243465).

## Supplementary Figure Legends

**Supplementary Figure 1:**
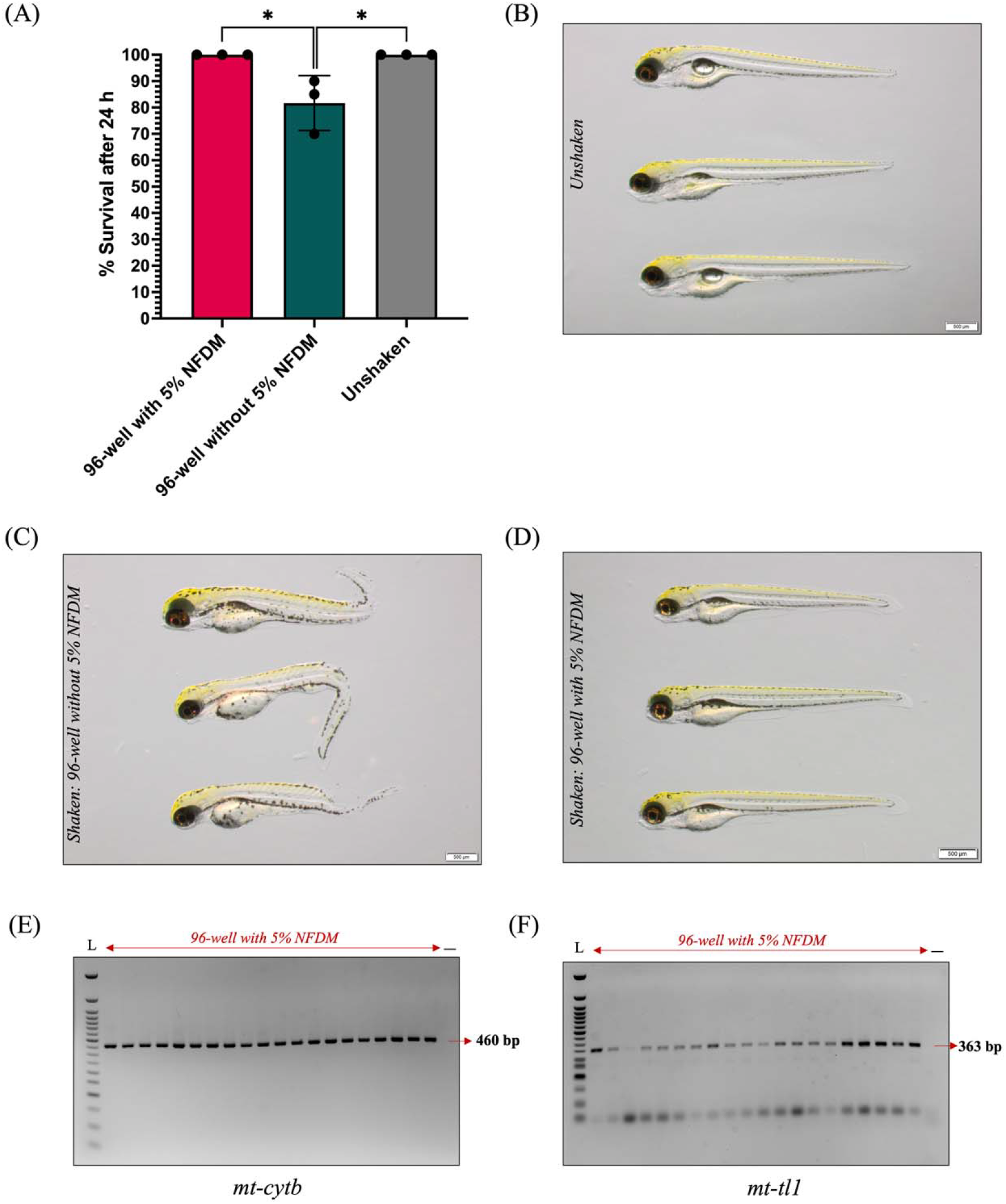
Effect of 5% non-fat dry milk (NFDM) solution on larval survival post genotyping. **(A)** Survival percentage of wild type larvae at 24 hours post genotyping in 96-well plate treated with NFDM, 96-well plate without 5% NFDM and unshaken control group. Bars represent survival percentage from three independent genotyping experiments. p-Values were determined by Student’s t-test (*p < 0.05; **p < 0.01; ***p<0.001). (N=60 technical replicates across 3 experiments). **(B-D)** Representative bright-field images of 3-dpf wild type larvae: **(B)** unshaken control, **(C)** 96-well plate without 5% NFDM showing tail damage in subset of larvae, and **(D)** 96-well plate with 5% NFDM showing undamaged, intact larvae. (Scale bar: 500 μm). **(E-F)** Gel electrophoresis demonstrating that 5% NFDM pre-treatment to 96-well plates does not interfere with the mitochondrial DNA genotyping, showing successful amplification of mitochondrial genes from larvae in 5% NFDM-treated 96-well plates: **(E)** *mt-cytb* (460 bp) and **(F)** *mt-tl1* (363 bp).

**Supplementary Figure 2:**
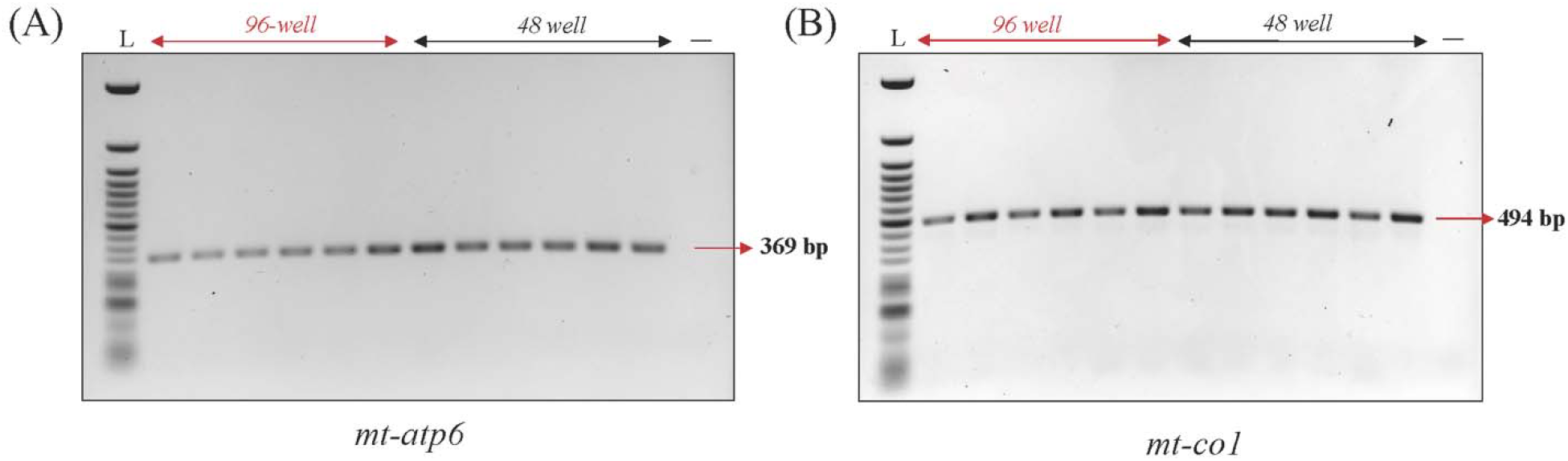
Mitochondrial DNA Amplification for zebrafish *mt-atp6* and *mt-co1*. **(A)** Gel electrophoresis of the mitochondrial gene *mt-atp6*, showing amplification of the 369 bp band. **(B)** Gel electrophoresis of the mitochondrial gene *mt-co1*, showing amplification of the 494 bp band.

**Supplementary Figure 3:**
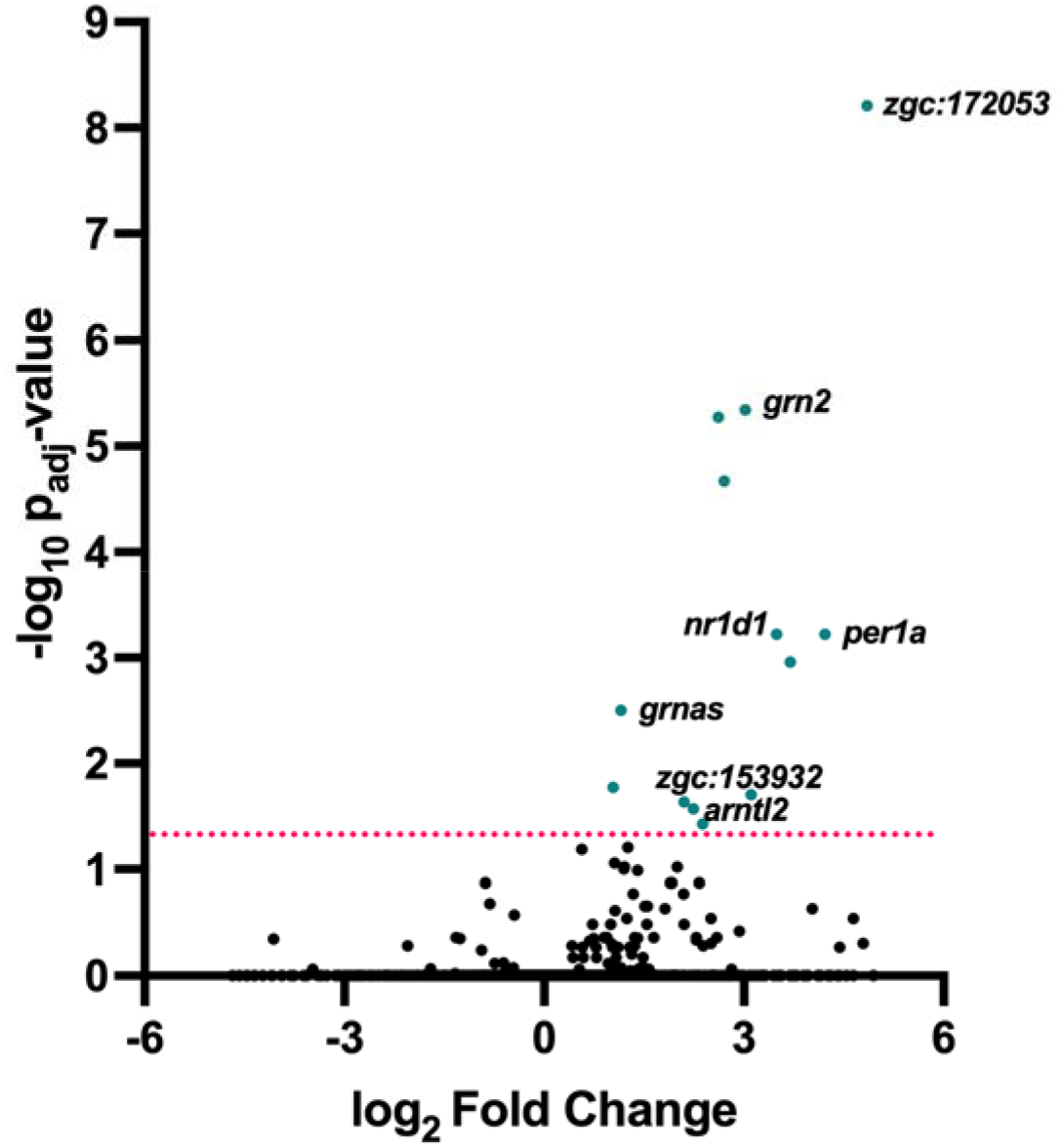
Volcano plot illustrating differentially expressed genes. The x-axis represents the log_2_ fold change in gene expression, with positive values indicating upregulation and negative values indicating downregulation. The y-axis shows the minus log_10_ of the adjusted p-value, representing statistical significance. Each dot represents a gene, with black dots indicating genes not significantly changed and green dots indicating significantly upregulated genes above the threshold (dotted line at adjusted p-value = 0.05).

## Supplementary Figure Legend

**Supplementary Table 1:**
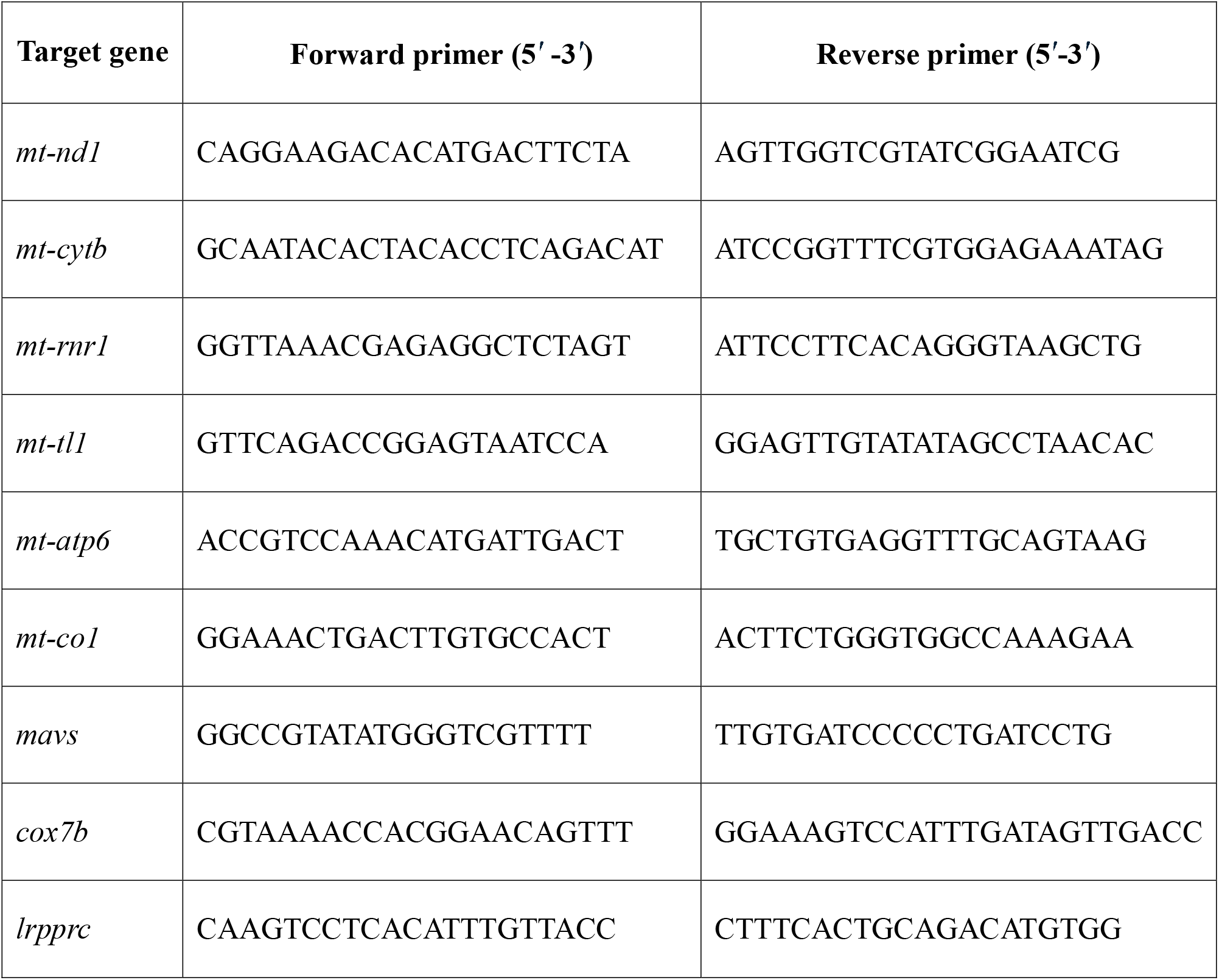
List of oligonucleotides used in this study.

## Supplementary File Legend

**Supplementary File 1: Fold change for all the genes from RNA sequencing experiment**

